# Crosslinker design determines microtubule network organization by opposing motors

**DOI:** 10.1101/2022.04.22.488007

**Authors:** Gil Henkin, Wei-Xiang Chew, François Nédélec, Thomas Surrey

## Abstract

During cell division, crosslinking motors determine the architecture of the spindle, a dynamic microtubule network that segregates the chromosomes. It is unclear how motors with opposite directionality coordinate to drive both contractile and extensile behaviors in the spindle. Particularly, the impact of different crosslinker designs on network self-organization is not understood, limiting our understanding of self-organizing structures in cells, but also our ability to engineer new active materials. Here, we use experiment and theory to examine active microtubule networks driven by mixtures of motors with opposite directionality and different crosslinker design. We find that although the kinesin-14 HSET causes network contraction when dominant, it can also assist the opposing kinesin-5 KIF11 to generate extensile networks. This bifunctionality results from HSET’s asymmetric design, distinct from symmetric KIF11. These findings expand the set of rules underlying patterning of active microtubule assemblies and allow a better understanding of motor cooperation in the spindle.

**SIGNIFICANCE STATEMENT:** During cell division, the spindle apparatus segregates duplicated chromosomes for their inheritance by the daughter cells. The spindle is a highly interconnected network of microtubule filaments that are crosslinked by different types of molecular motors. How the different motors cooperate to organize the spindle network is not understood. Here, we show that an asymmetric crosslinker design can confer bifunctionality to a mitotic motor in the presence of other motors. The asymmetric motor supports both extensile and contractile microtubule network behaviors as observed in different parts of the spindle. These findings define new rules controlling the generation of active microtubule networks and allow us to better understand how motors cooperate to organize the correct spindle architecture when a cell divides.

## INTRODUCTION

The mitotic spindle, which segregates the chromosomes during cell division, is a paradigm for self-organizing active filament networks. It is comprised of dynamic microtubules, interconnected by motile crosslinkers, forming an active gel continuously driven by internal stresses while maintaining steady-state shape^1, 2, 3, 4, 5^. The metaphase spindle can be conceptualized as a central extensile nematic network of antiparallel microtubules that gradually turns into polar microtubule networks at the spindle poles^6, 7^. How crosslinking motors arrange the microtubules in such a network is not understood.

In vitro reconstitutions of self-organizing microtubule networks with purified crosslinking motors have provided insight into the basic properties of active microtubule-based materials^5, 6, 8, 9, 10, 11, 12, 13, 14, 15, 16^. Extensile, nematic networks are promoted by high microtubule densities and growth rates, or the presence of crowding agents, whereas contractile or radially polar networks form when motors can accumulate at microtubule ends^6, 17^. The rules governing extensile versus contractile microtubule network formation when different crosslinking motors are present are, however, still unknown.

Moreover, natural motors have distinct designs. Kinesin-5 is a plus end directed, symmetric motor that can crosslink two microtubules using pairs of motor domains present at either end of the tetrameric molecule^18^. It slides antiparallel microtubules apart in the spindle center^8, 19, 20, 21^. Kinesin-14 is a minus end directed motor, thought to support pole focusing activity in conjunction with dynein^22, 23^. Unlike symmetric kinesin-5, kinesin-14 is an asymmetric motor that can crosslink two microtubules using one pair of motor domains and one pair of diffusible microtubule-binding domains at opposite ends of the dimeric molecule^8, 24, 25^. The consequences of different crosslinker design on network formation are unknown.

Here we perform in vitro experiments with mixtures of both mitotic kinesins and microtubules nucleating in solution. We find that crosslinker design plays a particularly important role when motors with opposite directionality work together to generate an active network. Kinesin-14, mostly known as an aster forming motor, promotes radially polar network formation when dominant, but is surprisingly also able to act cooperatively with kinesin-5 to generate nematic networks. Computer simulations demonstrate that this collective behavior is a consequence of kinesin-5 being a stronger crosslinking motor than kinesin-14 which on the other hand is still an efficient microtubule bundler. Our study illustrates how the design of mitotic motors is optimized for their combined action in the spindle and provides new insight into the principles of active multi-motor network organization.

## RESULTS

### The organizational capacities of HSET and KIF11 diverge at high tubulin concentrations

Recently, we systematically investigated the organizational phase space of microtubule self-organization driven by the symmetric crosslinker kinesin-5 ^6^. The capabilities of an asymmetric crosslinker, such as kinesin-14, to make diverse networks has not been studied as systematically yet. Using a microscopy-based motor/microtubule self-organization assay, we first assayed what network morphologies each type of crosslinkers could produce on its own (Fig. 1a), before studying how symmetric and asymmetric motile crosslinkers with opposite directionality work together. To promote efficient microtubule nucleation from soluble tubulin, we added low concentrations of the microtubule-stabilizing drug docetaxol, whose relatively high solubility compared to other taxanes was found to be advantageous (Methods). Under these conditions, microtubules polymerize until the tubulin subunits have been consumed and microtubules are then stable. Microtubule growth is intrinsically asymmetric, with plus ends growing faster than minus ends.

**Fig. 1.**
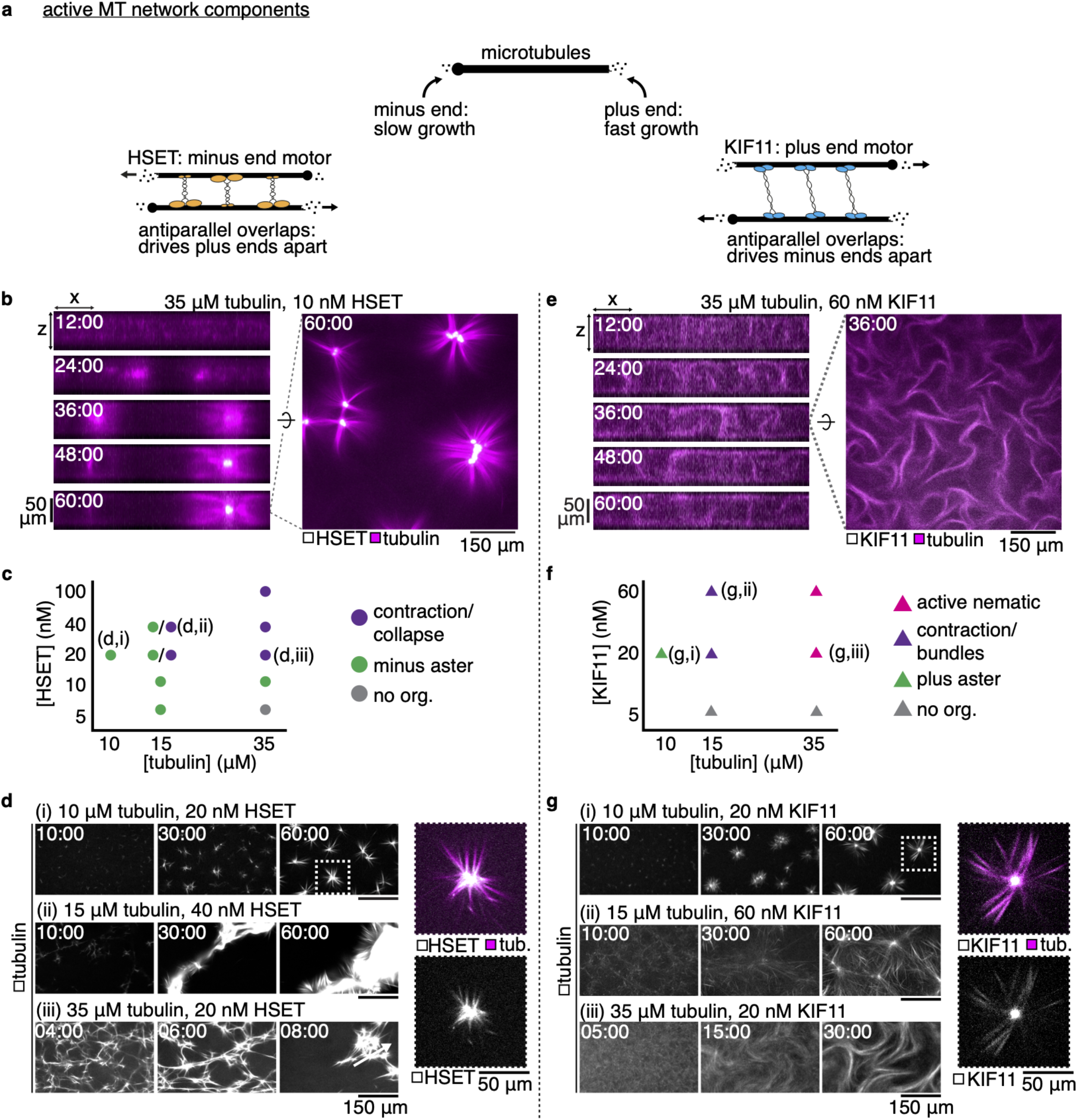
Characterizing HSET- and KIF11-driven organization of docetaxel-nucleated microtubule networks. **a**, Schematic showing the major components of competitive motor network assembly and their properties. **b**, Confocal imaging of the self-organization of a 35 μM tubulin network by 10 nM mCherry-HSET into separated asters. Left: Evolution of the network contraction into asters in an x-z plane. Right: Snapshot of HSET-mediated asters after 60 minutes of assembly in an x-y plane at the chamber’s midplane. Dashed line indicates the selected x-z slice. **c**, Phase diagram of mCherry-HSET-mediated microtubule network organization. **d**, Time sequence of microtubule organization for 3 selected conditions from the phase space, showing (i) minus asters, (ii) larger-scale contraction, and (iii) rapid network collapse. Right: Enlarged image of HSET aster; mCherry signal is limited to the pole-proximal microtubule lattice. **e**, Confocal imaging of the self-organization of a 35 μM tubulin network by 60 nM KIF11-mGFP into an active nematic network. Left: Evolution of the turbulent network extensile activity in an x-z plane. Right: Snapshot of KIF11-mediated extension after 36 minutes of assembly in an x-y plane at the chamber’s midplane. Dashed line indicates the selected x-z slice. **f**, Phase diagram of KIF11-mGFP-mediated microtubule network organization. **g**, Time sequence of microtubule organization for 3 selected conditions from the phase space, showing (i) plus asters, (ii) loosely contractile bundles, and (iii) active nematic activity. Right: Enlarged image of KIF11 aster; mGFP signal is highly enriched at the pole but still appears on distal microtubule lattice. See also Movie 1.

Purified minus-end directed human kinesin-14 HSET generated contractile microtubule networks as observed previously^6, 8, 10^. At relatively low tubulin and HSET concentrations (all motor concentrations refer to monomers), well-focused, separated asters formed (Fig. 1b). Asters became centered between the top and bottom of the experimental chamber and HSET accumulated in the middle of the asters, indicating that minus ends were gathered (Fig. 1b). Asters that came into contact fused (Movie 1a). We explored the phase space of network organization by HSET (Fig. 1c). Well-defined aster formation was favored at low tubulin concentrations (Fig. 1d i). At higher HSET and tubulin concentrations, when networks are expected to be more percolated, they contracted as large entities (Fig. 1d ii), as seen previously^6,8^. Contraction was fastest at high tubulin concentrations when microtubules nucleated and elongated at faster rates (Fig. 1d iii) (Suppl. Fig. 1). The overall contractile nature of HSET/microtubule networks is due to the relatively slow growth speed of microtubule minus ends compared to the HSET motor speed, allowing HSET to gather microtubule minus ends over a wide range of conditions^6^.

In contrast, purified plus-end directed human kinesin-5 KIF11 generated nematic networks of extensile bundles with evenly distributed KIF11 when the tubulin concentration was high (Fig. 1e, Movie 1b)^6^. The microtubule plus-end growth speed is high at early times under these conditions, which prevents KIF11 from gathering plus ends into contractile structures. Instead, KIF11 drives persistent sliding of antiparallel microtubules, which in the dense microtubule suspension leads to repeated buckling, merging and sliding of bundles in 3 dimensions (Fig. 1e)^6, 11, 14, 15, 16, 20^. We varied tubulin and KIF11 concentrations (Fig. 1f). At lower tubulin concentrations, at which microtubule plus ends grow more slowly, KIF11 formed contractile networks and individual asters, as it could now accumulate at microtubule plus ends and bring plus ends together (Fig. 1g i)^6, 13, 20^. Bundling and bundle extension was favored at higher tubulin concentrations (Fig. 1g ii-iii). These results agree with previous work where microtubules were nucleated differently^6^ and establish the organizational phase space for networks of docetaxel-nucleated microtubules organized by a single type of motile crosslinker, either HSET or KIF11.

The organizational capacity of the two motors at low tubulin concentrations is similar, but diverges at higher tubulin concentrations. This can be understood as a consequence of the opposite directionalities of the motors and the different growth speeds at the two microtubule ends. It is more difficult to for KIF11 to accumulate at the faster growing plus ends than HSET at the slower growing minus ends^6, 26^ when overall microtubule growth is faster.

### KIF11 antagonizes microtubule aster formation by HSET

Next, we set out to determine the network patterns formed in the presence of both motors.

First, we studied low tubulin concentration networks under conditions where each motor individually would drive contractile network behavior or generate asters, however with opposite polarity. When KIF11 was present at a fourfold lower concentration than HSET, KIF11 delayed HSET-driven aster formation (Fig. 2a i, Movie 2a). Initially, microtubules appeared heavily crosslinked into dense clusters without obvious orientation or polarity. Over time, these clusters reorganized into asters with central HSET accumulation, revealing that when in excess HSET can out-compete KIF11 under aster-promoting conditions (Fig 2a ii). Network organization took more time with both motors being present than with each motor individually, suggestive of a sustained competition for dominance. The presence of KIF11 between HSET-enriched centers prevented these polar structures from fusing, despite persistent contact (Fig. 2a i).

**Fig. 2.**
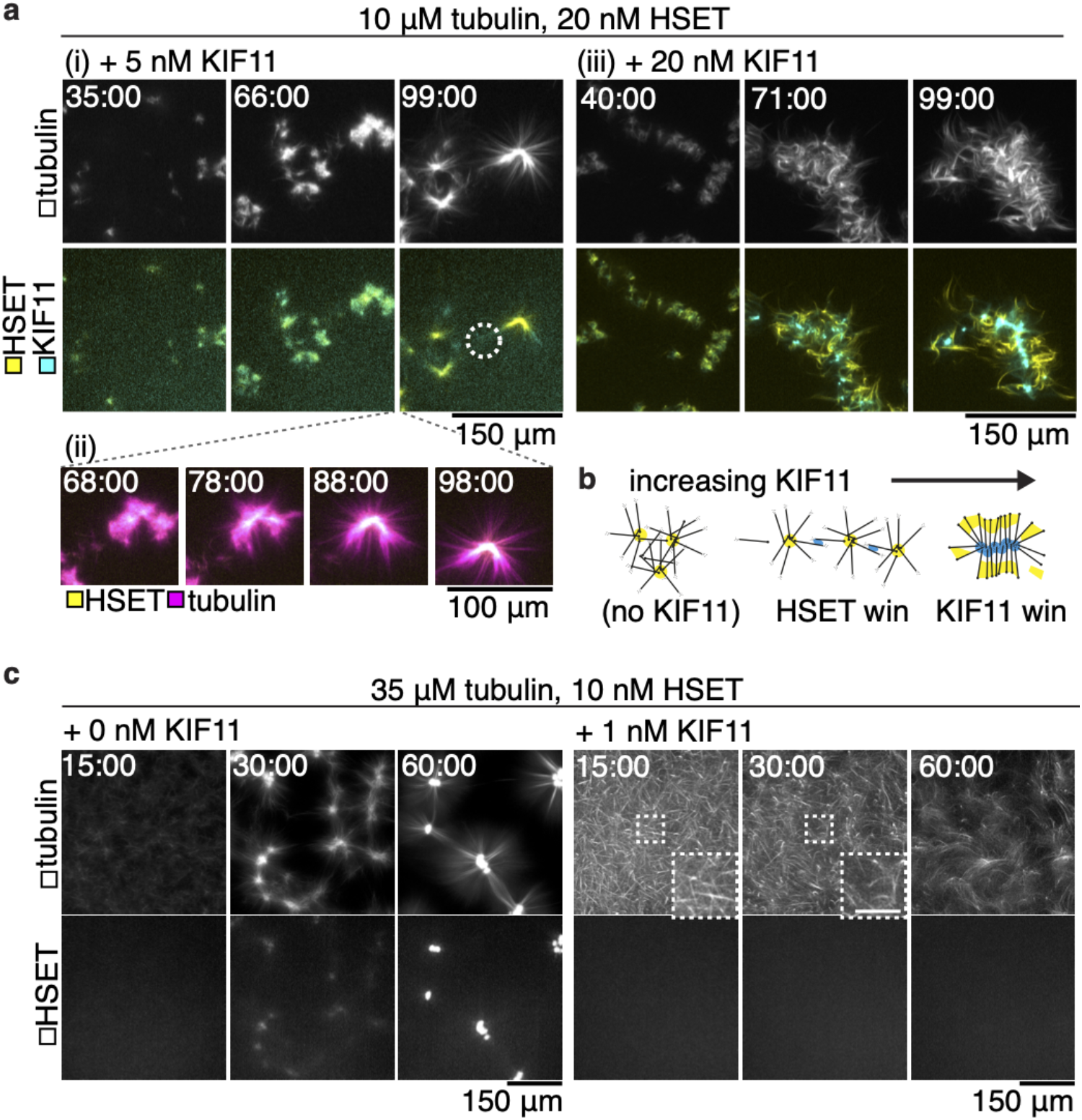
HSET and KIF11 compete to organize microtubule networks. Time courses show maximum projections of confocal images taken around the chamber midplane. **a**, mCherry-HSET and KIF11-mGFP (bottom panels) compete to contract low-density microtubule networks (top panels). (i) 5 nM KIF11-mGFP frustrates assembly of asters in 10 μM tubulin networks (compare to **Fig. 1c**) and eventually appears as a diffuse signal decorating the microtubules between mCherry-HSET contractile centers (circle). (ii) Localization of HSET within the microtubule structure coincides with the radial appearance of microtubule ends. (iii) 20 nM KIF11-mGFP dominates assembly of locally-contractile structures and localizes to the structure centers. **b**, Schematic of competitive assembly of contractile microtubule networks by KIF11-mGFP and mCherry-HSET. **c**, Left, 10 nM mCherry-HSET (bottom panels) contracts microtubules (top panels) assembled from 35 uM tubulin into asters with well-defined centers (as in **Fig. 1b**). Right, the addition of 1 nM KIF11-mGFP prevents HSET contraction. mCherry-HSET signal remains diffuse within the microtubule network. Microtubule bundles (insets, scale bar 50 µm) increase in curvature, indicating strain within the network. See also Movie 2.

At equimolar monomer concentrations, the two motors again formed densely crosslinked microtubule clusters which coalesced as they came into contact. The motors subsequently self-sorted (Fig. 2a iii, Movie 2b). KIF11 resided at the center of the assembled structures, indicating that it had successfully gathered the plus ends and dominated the contractile network behavior. HSET was relegated to the periphery where it bundled microtubules, but did not appear to efficiently gather minus ends (Fig. 2a iii). Thus, in the low tubulin, contractile regime, the motor ratio determines which motor wins the competition (Fig. 2b), with KIF11 being the more efficient competitor.

The dominance of KIF11 in driving network behavior was even more striking at high tubulin concentrations. At a concentration of HSET that causes the formation of well-defined, large asters in the absence of KIF11 (Fig. 2c left), addition of a tenfold lower concentration of KIF11 completely inhibited aster formation or any local accumulation of HSET (Fig. 2c right). Instead, a crosslinked mesh of curved microtubule bundles formed, characteristic of extensile network behavior. These results show that in dense networks with higher microtubule growth rates, in which HSET forms asters only within a narrow range of concentrations (Fig. 1c), KIF11 is a particularly efficient competitor for HSET.

### Tubulin concentration is a control parameter influencing nematic versus radially polar organization in mixed-motor networks

We saw a marked difference in mixed motor network organization outcomes between low and high tubulin concentrations. To investigate the effect of microtubule growth speed and density on network formation in the presence of the two motors, we allowed networks to organize at increasing tubulin concentrations while keeping the HSET and KIF11 concentration constant, at levels where KIF11 is dominant (Fig. 3). At low tubulin concentrations, contractile networks formed, and KIF11 eventually localized to astral centers (Fig. 3a, same conditions as in Fig. 2a ii). Increasing the tubulin concentration led to more long-range microtubule bundling and less local contraction (Fig. 3b, c). Motor self-sorting to different bundles of the network was increasingly hindered with increasing tubulin concentrations. At the highest tubulin concentration tested, the network was mostly nematic displaying extending bundles and evenly distributed motors (Fig. 3d).

**Fig. 3.**
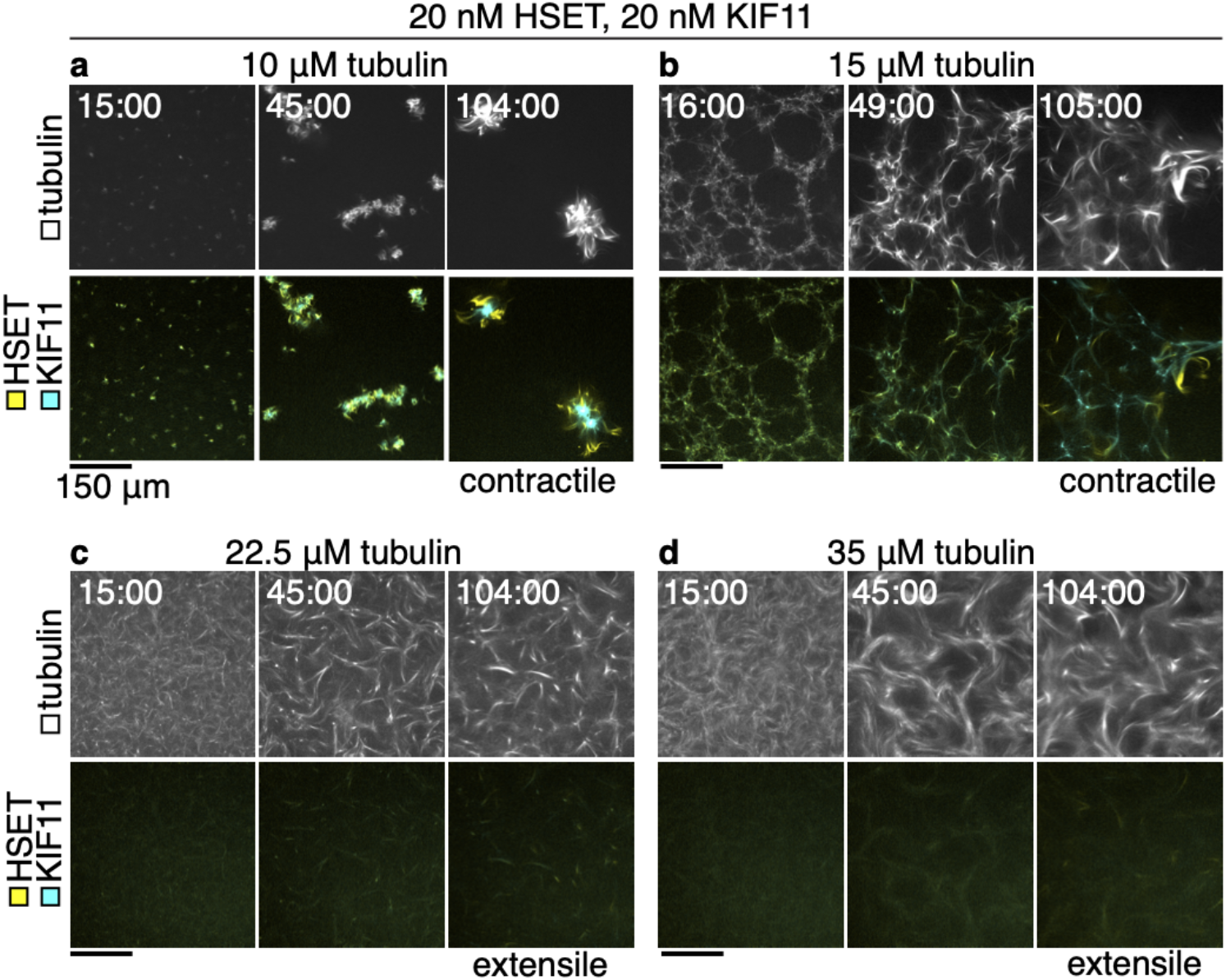
Increasing tubulin concentration drives transition from contractile to extensile gels in competitive motor-driven networks. Time course of confocal images taken in the flow-cell midplane of microtubules (top panels) organized by 20 nM mCherry-HSET and 20 nM KIF11-mGFP (bottom panels). **a**, As in **Fig. 2a**, microtubules polymerized from 10 μM tubulin are contracted and form compact, locally connected structures. KIF11 is localised to the structure center, and HSET is excluded to the periphery. **b**, 15 μM tubulin networks form contractile webs with long-range interactions. KIF11 signal is internal to the contracting web, HSET localizes to bundles extruding from the network. **c**, 22.5 μM tubulin assembles into microtubules that form extensile bundles. Motors are well-distributed in the bundles, but HSET localization at bundle tips can be seen emerging from the extensile network. **d**, 35 μM tubulin assembles into microtubules that form extensile bundles. Differential localization of the motors within the same time-frame is less clear. All scale bars 150 μm.

In conclusion, increasing microtubule density and growth speed by increasing the tubulin concentration changed the nature of the ‘mixed motor’ network from contractile to nematic, similar to the trend in ‘KIF11 only’ networks (Fig. 1e). HSET could accumulate locally in bundles but could not efficiently generate focused minus ends. HSET delayed, but did not prevent, KIF11-driven network organization, indicating that HSET plays an antagonistic role when KIF11 dominates.

### HSET supports KIF11 in driving nematic network formation at low KIF11 concentration, revealing multi-functionality

Next, we investigated more closely the high-tubulin concentration regime, in which the independent organizational capacities of the two motors diverge, with HSET forming contractile and KIF11 extensile networks (Fig. 1). Keeping the tubulin concentration constant, we added increasing concentrations of HSET to KIF11 kept at a concentration that was too low to organize a macroscopic network state (Fig. 4a i, Movie 3, top left). We found that addition of an equimolar amount of HSET resulted in weak nematic network formation with some extensile activity (Fig. 4a ii, Movie 3, top right). Increasing the HSET concentration 2-fold and 8-fold produced increasingly active nematic networks with long, dense microtubule bundles (Fig. 4a iii,iv, Movie 3, bottom left). This is remarkable, because HSET on its own is unable to promote nematic network formation in these experimental conditions (Fig. 1c). HSET distributed throughout the bundles at an 8-fold excess of HSET over KIF11 (Fig. 4a iv). Increasing the HSET concentration further to a 20-fold excess over KIF11 finally caused the network to become contractile; initially microtubules assembled into bundles that eventually reformed into aster-like structures (Fig. 4a v, Movie 3, bottom right).

**Fig. 4.**
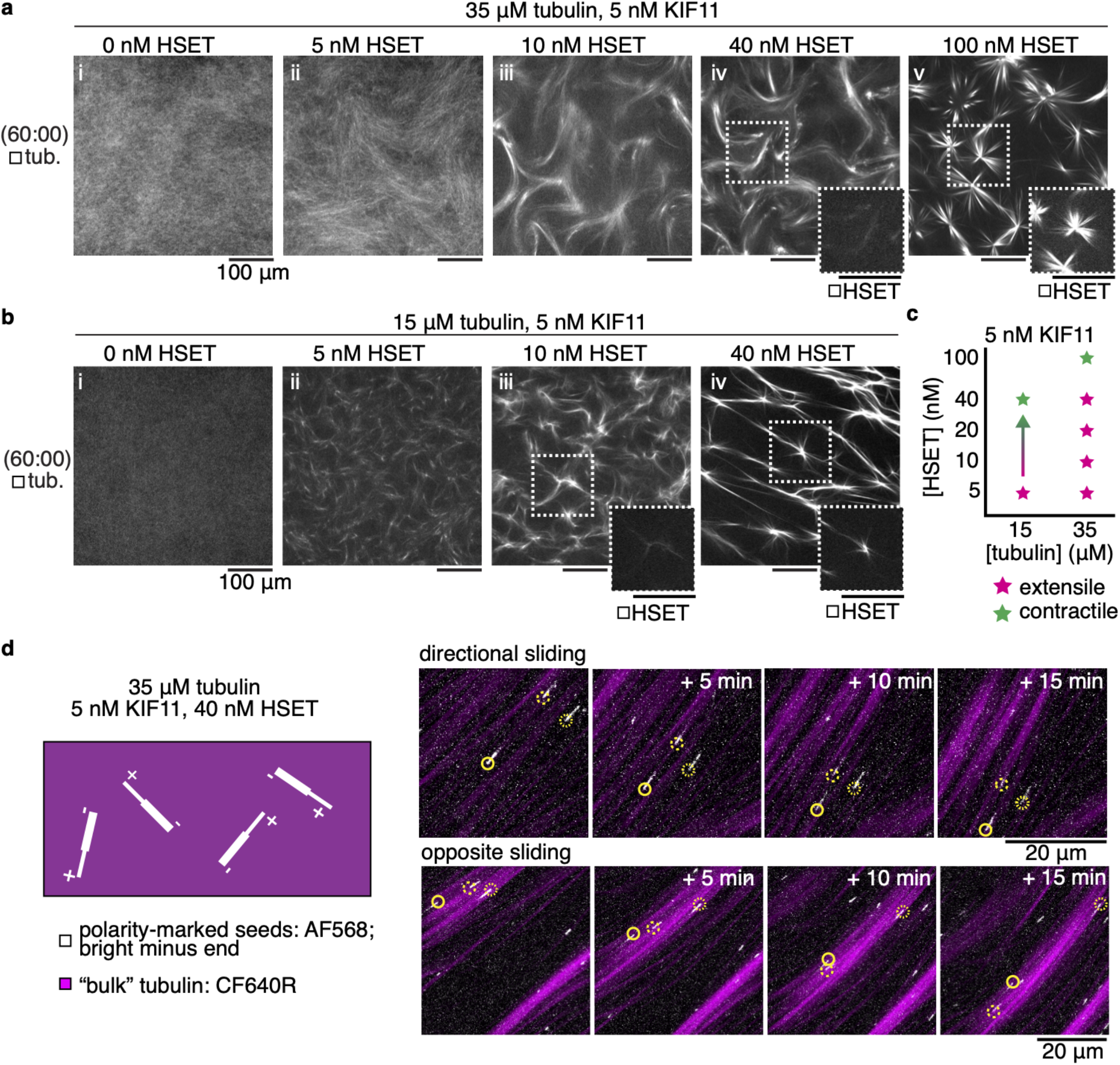
HSET facilitates extensile behavior in mixed-motor networks consistent with plus-end directionality. **a, b**, Snapshots show midplane confocal slices of the experimental chamber after 60 minutes of self-assembly. **A**, Addition of (i) 5 to (iv) 40 nM mCherry-HSET drives extensile behavior of 35 μM tubulin networks in the presence of 5 nM KIF11-mGFP, which is insufficient to drive visible organization on its own. (v) 100 nM HSET drives contractile organization. HSET localizes to extensile bundles and contractile centers (insets). All scale bars 100 µm. **b** Addition of (ii) 5 nM HSET drives extension of 15 uM tubulin in the presence of 5 nM KIF11 whereas 15 μM tubulin networks with either motor alone only show contractile behavior (see **Fig. 1c, f**), or, as in the case of (i) 5 nM KIF11, no organization. (iii) 10 to (iv) 40 nM HSET shows tendency towards contraction of the network, where mCherry-HSET localizes to contractile centers (insets). All scale bars 100 µm. **c**, Results summarized in a phase space plot. **d**, Time courses show cropped maximum-projections of confocal images taken in thin (20 μm) chambers. A small number of seeds with bright minus ends (AlexaFluor567) demonstrate individual microtubule activity in a background of tubulin with a spectrally-separate fluorescent dye (CF640R). Arrows indicate the direction of the microtubule plus end. Microtubules in networks assembled from 35 μM tubulin, 5 nM KIF11-mGFP, and 40 nM mBFP-HSET generally move towards their minus ends (top panels) and could be seen moving in opposite directions within aligned bundles (bottom panels). See also Movies 3-5.

Interestingly, although the motors have opposite directionality, HSET can support KIF11 to drive the formation of extensile bundles even at lower tubulin concentrations (Fig. 4b, Movie 5) which otherwise promote contractile activity with either motor alone (Fig. 1c, f). The range of HSET concentrations that assist extensile bundle formation driven by KIF11 is however decreased compared to high tubulin concentration networks (Fig. 4b i-iii). This is a consequence of contractile behavior being recovered more easily in these sparser networks with fewer and slower-growing microtubules, and aster-like structures are recovered already at an 8-fold excess of HSET over KIF11 (Fig. 4b iv, Fig. 4c). Motors probably accumulate more efficiently at microtubule ends, which then promotes network contraction and aster formation. The asymmetric crosslinker HSET may assist the symmetric crosslinker KIF11 to drive nematic network formation of extensile bundles by providing additional microtubule crosslinks despite losing the competition for directional sliding. This is likely due to the weaker processivity of the HSET motor and its diffusive microtubule binding domain that prevents strong force production, which would mean that KIF11 drives microtubule sliding in these mixed motor nematic networks^24, 27, 28^. To test this hypothesis, we added polarity marked seeds to high-tubulin concentration networks with an 8-fold excess of HSET over KIF11 and found that the majority of seeds were indeed moving with their minus end leading within the bundles (Fig. 4d, Suppl. Fig. 2, Movie 5), indicating KIF11-driven microtubule sliding. Some seeds displayed saltatory motility, indicative of a “tug of war” in some parts of the network^8, 29^, although the overall macroscopic dynamics were dominated by KIF11.

These experiments demonstrate that the asymmetric design of HSET, with one motor and one diffusible microtubule binding domain, turns this motor into a multifunctional crosslinker that can either assist KIF11 in nematic network formation or promote contractile network and aster formation depending on the relative motor concentrations.

### Computer simulations with symmetric and asymmetric crosslinkers recapitulate experimental network organization

To pursue our investigation of these mixed motor microtubule networks, we turned to computer simulations. Microtubules were modelled in a thin three-dimensional space as diffusing bendable lines repelling each other via soft-core interactions^30, 31^. They grew from a fixed number of nucleators by plus-end elongation for a defined period of time and then kept a fixed length (Suppl. Fig. 3a), mimicking the behavior of the docetaxol-stabilized microtubules in the experiments. KIF11 was modelled as a symmetric complex consisting of two plus-end directed and processive motor units mechanically connected by a Hookean spring with non-zero resting length (Fig. 5a)^30^. KIF11 units unbound immediately upon reaching microtubule plus ends (see Methods). In contrast to previous microtubule network simulations, we explicitly considered here the asymmetric nature of HSET. It was modelled as an asymmetric crosslinker with one minus-end directed, non-processive motor unit and one diffusive unit (Fig. 5a). The dwell time of the diffusive unit was 500 times larger than that of the motor unit, to reflect the experimentally measured properties of this motor (see Table S1 for model details)^24^. Both HSET units dwell at microtubule ends, not detaching immediately (see Methods).

**Fig. 5.**
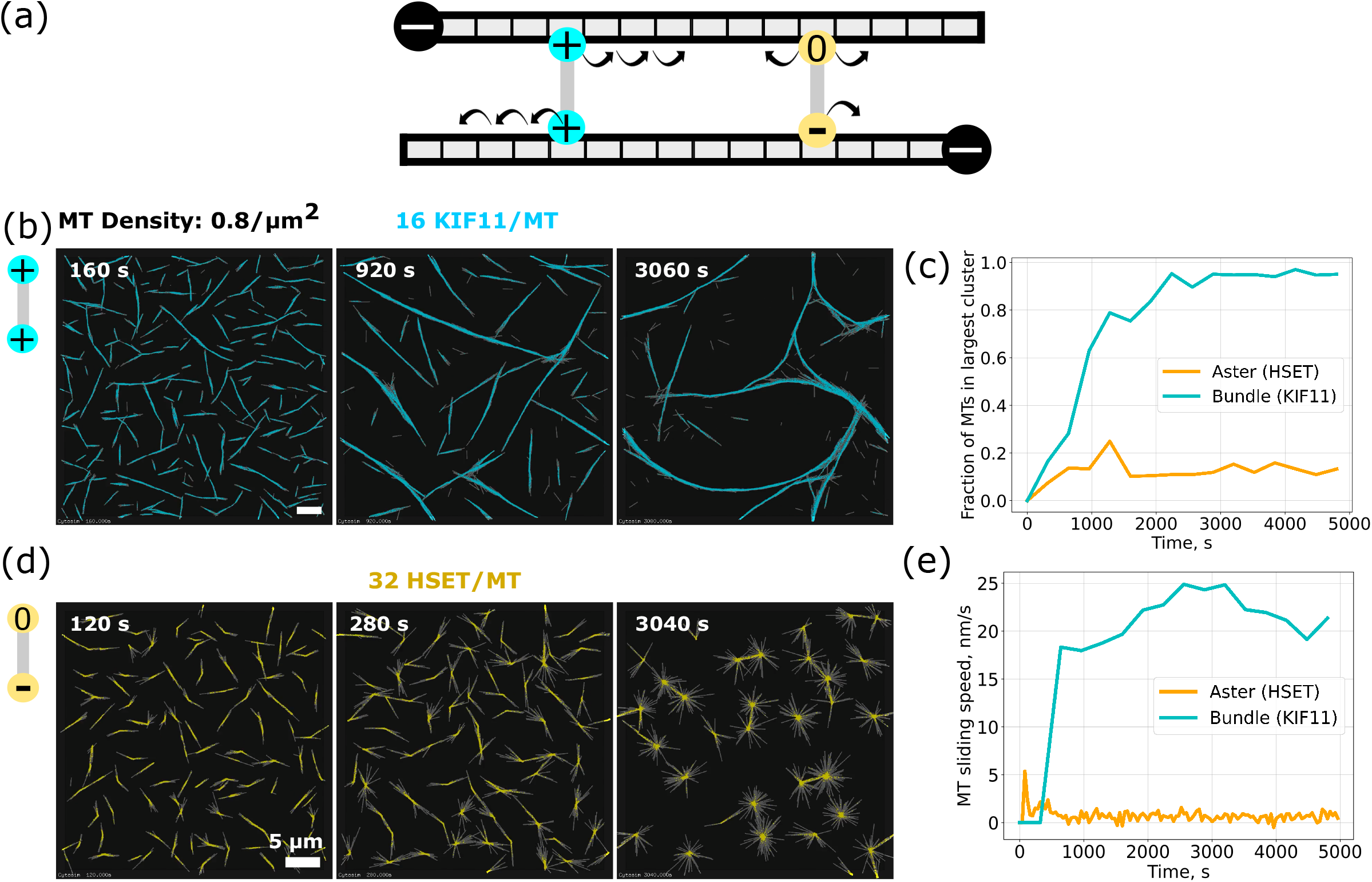
Simulated KIF11 and HSET-driven microtubule organization. **a**, Model schematic of KIF11 (cyan) and HSET (yellow) crosslinker. KIF11 is a symmetrical crosslinker composed of two plus-end directed motor units connected by a spring-like link. HSET is an asymmetrical crosslinker made of one minus-end directed motor unit and one diffusive unit. **b**, Time course of simulated KIF11-driven extensile nematic bundle in a thin 3D box (dimension: 60×60×0.2 µm). The 2880 microtubules are colored in gray, and the 46080 KIF11 in cyan. **c**, Time series of largest bundle size (cyan line) and aster size (orange line) from the same simulations. Sizes were normalized by the number of microtubules. **d**, Time course of simulated HSET-driven aster formation. Simulation space is 40×40×0.2 µm with 1280 microtubules and 40960 HSET. Scale bar is 5 µm. Model parameters are given in (Table S1). **e**, Displacement speed of microtubules in KIF11-driven (cyan line) and HSET-driven (orange line) organization. Total simulation duration is 5000 s. See also Movie 6.

We first examined networks driven solely by HSET and KIF11. In the absence of any motor, microtubules remained randomly oriented (Suppl. Fig. 3b). At a microtubule density of 0.8 microtubules/µm^2^, KIF11 organized microtubules into bundles that continuously extended, broke apart and reformed again (Fig. 5b, Movie 6a), similar to experimentally observed active nematic networks driven by KIF11 at high tubulin concentrations (Fig. 1e). To characterize the network, we extracted key quantities from the simulations: the number of crosslinkers in each configuration, the number of microtubules in a cluster and the speed at which microtubule moved (Methods). In KIF11-driven networks, the average microtubule sliding speed increased with the number of microtubules becoming incorporated in the mixed polarity bundles (Fig. 5c, e, cyan). In contrast, HSET bundled and organized microtubules into asters with focused minus ends (Fig. 5d, Movie 6b) leading to immobile microtubules (Fig. 5c, e, orange), similar to experimentally observed asters generated by HSET (Fig. 1b). This shows that an asymmetric crosslinker with one diffusive and one motor unit, even lacking processivity, can form asters, without requiring larger motor oligomer formation^10^. Compared to simulations with a hypothetical symmetric minus-end directed motor, made of two mildly processive motor units, the modelled HSET crosslinker drove assembly of asters with less focused centers, and caused microtubules to be more bundled close to the aster center (Suppl. Fig. 4, left and middle), similar to experimentally observed HSET asters (Fig. 1d i, inset), further validating the HSET model.

### HSET activity synergizes with or antagonizes KIF11 depending on relative abundance

To determine why and under which conditions HSET either antagonizes or cooperates with KIF11, we simulated microtubules with few KIF11 crosslinkers, insufficient to organize microtubules (Fig. 6a left, Movie 7a). Adding a low amount of HSET crosslinkers promoted bundle formation (Fig. 6a middle, Movie 7b). Visualizing only some of the microtubules in the simulated bundles revealed mostly KIF11-driven microtubule sliding and occasionally saltatory movement, resembling experimental observations (Fig. 6b, Movie 8)^8, 29^. These simulations demonstrate that, as in the experiments (Fig. 4), despite its opposite directionality, low amounts of HSET assist KIF11-driven extensile bundle formation. That is because HSET can contribute efficiently to bundling, even though it can only produce weak microtubule sliding forces (Suppl. Fig. 5) as a consequence of the diffusive binding mode of its tail^27^. Nevertheless, the average microtubule sliding speed in these mixed motor bundles is slower than in ‘KIF11 only’ bundles, indicating that HSET crosslinking does generate an opposing frictional force slowing down KIF11-driven sliding (Fig. 6c). Further increasing the amount of HSET reduces the microtubule sliding speed (Fig. 6c) and at even higher amounts leads to HSET-driven aster formation, with microtubules becoming immobilized by anchoring of their minus ends in the aster center (Fig. 6a right, c, Movie 7c), again similar to the experiments (Fig. 4a). Thus, whereas at lower amounts HSET assists KIF11, at the high amounts HSET antagonizes and outcompetes KIF11.

**Fig. 6.**
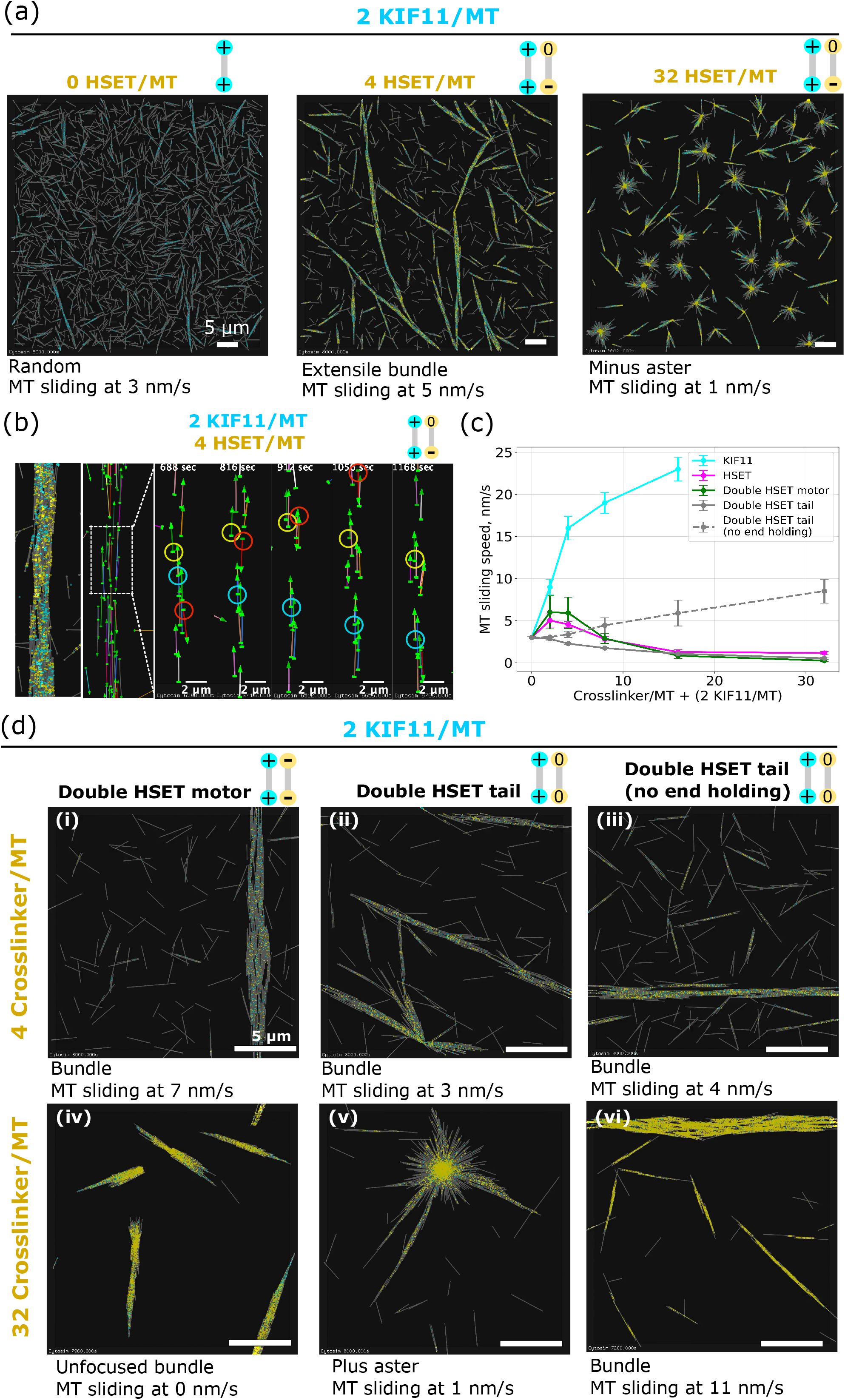
Simulated microtubule network organized by mixed motors. **a**, Mixing a small amount of KIF11 with an increasing amount of HSET. Left: Random microtubule organization in the presence of low amount of KIF11 (cyan); Middle: Extensile bundle formed when HSET (yellow) was added in the presence of KIF11 at 2:1 ratio. Right: Minus aster formation when HSET and KIF11 were mixed at 16:1 ratio. Simulation space is 60×60×0.2 µm with 2280 Microtubules and 5750 KIF11; Total simulation duration is 8000 s. See also Movie 7. **b**, Visualization of selected individual microtubule trajectories in mixed motor condition (2 HSET/KIF11). Majority of microtubules, depicted as arrows, move unidirectionally with the minus end leading, sign of KIF11-driven sliding. See for example red circles. Back-and-forth motion occur occasionally (yellow circles). See also Movie 8. **c**, Overall microtubule sliding speed as a function of crosslinker number for various types of crosslinker mixed with a fixed amount of KIF11 (2 KIF11/MT). Each point represents speed averaged from 5 samples simulation run in a box of size 20×20×0.2 µm. d, Microtubule network formed by mixing variant of HSET crosslinker (yellow) with fixed amount of KIF11 (cyan) at 2 KIF11/MT. Variant crosslinkers are made of two HSET motors (i & iv), two HSET tails (ii & v) and two HSET tails but without microtubule end holding (iii & vi). Top row: low amount of variant crosslinker (4 crosslinkers/MT); Bottom row: high amount of variant crosslinker (32 crosslinkers/MT). Simulation space is 20×20×0.2 µm with 320 microtubules, covering 8000 s in total. All simulation contains the same density: 0.8 microtubule /µm^2^.

### The asymmetric design of HSET explains its multifunctional nature

To understand which properties of HSET makes it able to either synergize with or antagonize KIF11, we simulated variations of the HSET model, combining its different subunits into symmetric constructs. Symmetric crosslinkers consisting of two identical HSET motor units (‘double-motor HSET’) were equally efficient as the normal HSET in helping KIF11 to organize microtubules into bundles (Fig. 6d, i). The bundles extended, driven by KIF11, as long as a low amount of double-motor HSET was present (Fig. 6c). Higher amounts of double-motor HSET promoted stronger bundle formation, but failed to generate asters (Fig. 6d, iv). Instead, microtubules were polarity sorted into bundles with double-motor HSET accumulating weakly at the microtubule ends. This demonstrates that the presence of the long-dwelling HSET tail is important for HSET’s ability to form asters. The long dwell time of the HSET tail allows its non-processive motor to quickly rebind to a microtubule when tethered in a bundle, effectively increasing the overall processivity^32^, and hence its ability to efficiently gather microtubule minus ends.

Adding low amounts of another symmetric crosslinker variant, now consisting of two diffusive HSET units (‘double-tail HSET’), also helped KIF11 to form microtubules bundles (Fig. 6d, ii). Bundling was more efficient than with normal HSET, given the long dwell time of the two diffusive units. However, KIF11-driven bundle extension was slower as a consequence of the increased friction produced by the diffusive units. Surprisingly, at higher amounts of double-tail HSET, a qualitatively different network formed: KIF11-driven asters with plus-end poles (Fig. 6d, v). In these asters, long-lived double-tail HSET crosslinks could keep plus ends of microtubules together after they were slid apart by KIF11 (Suppl. Fig. 4 right). KIF11-driven plus-end aster stabilization by double-tail HSET required the ability of the diffusive unit to hold on to microtubule ends. When the diffusive units in the model were allowed to unbind from microtubule ends, KIF11-driven extensile bundles formed instead of plus-end asters (Fig. 6d, iii, vi).

These simulations demonstrate how important the specific properties of crosslinkers are for their propensity to support the formation of different types of active networks, especially when multiple motor types are active. The design of the asymmetric crosslinker HSET combines properties that favor context-dependent microtubule bundling or aster formation. This unique set of properties make HSET multifunctional, supporting either plus motor-driven nematic network formation when present in lower numbers or minus-end aster formation when dominant.

## DISCUSSION

We observed both contractile and extensile network states in active microtubule networks generated by motors with opposite directionality, depending on system composition. Due to the asymmetric design of the minus motor HSET and the symmetric design of the plus motor KIF11, the two motors behaved differently. They do compete, but especially in the higher microtubule density regime, HSET can also assist KIF11 in nematic network formation, effectively creating a crosslinker with two functionalities depending on context.

HSET’s asymmetric design, characterized by a non-processive, minus-end directed motor that is diffusively anchored to a second microtubule for relatively long times, allows efficient aster formation in the absence of other motors, although it does not focus microtubules minus ends into very tight poles. Against an opposing motile crosslinker like KIF11 with two processive motors, HSET’s weak force producing capacity as a consequence of its tail’s diffusive binding^27^ requires a large excess over KIF11 to be able to compete efficiently and to impose minus aster formation. At similar concentrations of the two motors, new behavior emerges: HSET contributes to microtubule bundling and subsequently loses the competition for microtubule sliding, now effectively assisting KIF11-driven nematic network formation, slowing down microtubule sliding only mildly.

We previously demonstrated how the characteristically asymmetric properties of microtubule growth affect the generation of active networks by a single type of motor. Fast growth of microtubule plus ends favors nematic network formation by plus end directed crosslinkers, whereas slow (or no) minus end growth favors the formation of asters with a minus pole by minus end directed crosslinkers^6^. This raised the interesting question which network types dominate when opposite directionality motors are mixed, going beyond previous studies where only aster regimes were accessible^8, 12^. We showed here that the emergent phenomena in mixed-motor networks can be explained by the crosslinking structures of the motors, and not only by the simple rules that determine organization by single motor species.

The specific design of these two motors appears to be optimized for their function in the mitotic spindle. HSET is distributed over the entire spindle, in agreement with a role for microtubule bundling, indicating that it may assist KIF11-driven sliding of microtubules towards the poles also in cells. This assisting role may explain why HSET levels contribute to controlling spindle length in a previously counter-intuitive manner^33^. Kinesin-14 however also promotes pole focusing and bipolarization in cancer cells with too many centrosomes^34, 35^ and in meiotic oocytes lacking centrosomes^36, 37, 38^, supporting the notion that its function can differ in a spindle region where microtubule minus ends are enriched, cooperating there with the minus motor dynein.

Biological components with designs optimized for different tasks in living cells offer a variety of “tools” for integration into biomimetic materials, enriching the rulebook by which we understand macroscopic self-organized states, while giving rise to surprising new behaviors^14, 15, 16^. Here, we have shown how motile crosslinkers determine the large-scale properties of an active material and how these properties can be controlled not only by varying crosslinker concentrations, but also crosslinker properties. Despite our systematic exploration, we did not observe nematic and polar networks coexisting as in mitotic spindles. This most likely indicates that additional components are still missing to form spindle-like bipolar structures. Indeed, fine local control of microtubule nucleation and dynamics are well-known parameters of mitotic spindles that still need to be added *in vitro* for the active network to self-organize into novel patterns. This is an important challenge for bottom-up assembly of active microtubule assemblies in the future.

## METHODS

### I. EXPERIMENTS

#### Tubulin

Tubulin was initially purified by two rounds of polymerization and depolymerization from porcine brain as described previously^39^. N-ethylmaleimide (NEM) tubulin was prepared from purified tubulin as described previously^40^ without further recycling steps. To obtain small aliquots of highly active tubulin, further recycling and labelling with CF640R-NHS or AlexaFluor568-NHS was performed as described previously^40^. Final concentrations and labelling ratios were determined by nanodrop (extinction coefficient 115,000 M^-1^cm^-1^, molecular weight 110 kDa) before adjustment, final ultracentrifugation, and aliquoting (unlabeled tubulin: 15 μL aliquots, 200 μM; labelled tubulin: 2 μL aliquots, 150 μM). Aliquoted tubulin was then snap-frozen and stored in liquid nitrogen.

#### Recombinant motor proteins

Insect cell expression constructs for StrepTagII-mCherry-G_5_A-HSET (pJR291) and StrepTagII-KIF11-A_3_G_5_-mGFP (pJR303) have been described previously^6^. To generate an N-terminal mBFP-tagged HSET construct, similarly to previously described biotinylated constructs^41^, HSET was cloned into a modified pFastBacDual vector containing a biotin-acceptor peptide (BAP), mBFP, and flexible linkers, resulting in the expressed construct BAP-G_5_-mBFP-ELG_6_A-HSET, which co-expresses with the biotin ligase BirA from separate promoters (pJR349). This construct was used for polar seed assays as a spectrally different alternative to mCherry-HSET – the biotin tag was unused and did not interfere with HSET activity.

Recombinant motor proteins were expressed in cultures of Sf21 insect cells according to the manufacturer’s protocols, cell pellets from 500 mLs of culture were snap-frozen, and stored at -80°C. mCherry-HSET and KIF11-mGFP were purified by affinity chromatography using Strep-Tactin resin as described previously^6^. StrepTagII domains were cleaved by TEV protease after elution, and cleaved protein was further purified by gel-filtration. KIF11-mGFP was supplemented with sucrose to 10% w/v before the final ultracentrifugation, and both proteins were snap-frozen in 5 μL aliquots at 1 mg/mL and stored in liquid nitrogen (Fig. S5a, b).

Cell pellets expressing BAP-mBFP-HSET and BirA were resuspended in ice-cold lysis buffer (50 mM Na-phosphate, 300 mM KCl, 5mM MgCl_2_, 1 mM EGTA, 5 mM β-mercaptoethanol, 0.5 mM ATP, pH 7.5), supplemented with DNAseI and protease inhibitors. Lysate was homogenized in a glass douncer on ice, clarified by centrifugation at 50,000 RPM in a Ti70 rotor at 4°C for 45 minutes, and filtered using disposable syringe filters with a 0.45 μm pore size. Biotin from the media was removed by buffer exchange over HiPrep desalting columns equilibrated in lysis buffer. The lysate was loaded onto immobilized monomeric avidin beads, and bound protein was washed with lysis buffer, HSET gel-filtration buffer (50 mM Na-phosphate, 300 mM KCl, 1 mM MgCl_2_, 1 mM EGTA, 5 mM β-mercaptoethanol, 0.1 mM ATP, pH 7.5) supplemented with 5 mM ATP, and finally HSET gel-filtration buffer. After eluting protein with HSET gel-filtration buffer supplemented with 5 mM biotin, biotin was again removed by running over a HiPrep desalting column equilibrated in HSET gel-filtration buffer. The protein was gel-filtered using a Superose 6 10/300 GL column, concentrated to 1 mg/mL, clarified in a table-top ultra-centrifuge, snap-frozen in 5 μL aliquots, and stored in liquid nitrogen (Fig. S5c).

Final recombinant protein concentrations were determined by Bradford assay against a BSA standard on snap-frozen and thawed protein aliquots. Reported concentrations refer to monomers based on their predicted molecular weights.

#### Glass

Polyethylene glycol (PEG)-passivated glass was prepared as described previously^42^, without inclusion of biotinylated PEG-silane. Briefly, piranha-cleaned coverslips were thoroughly dried and incubated with GOPTS by making “sandwiches” in petri dishes on a 75°C hot plate, pressing out air bubbles, followed by incubation in a 75°C oven. After 30 minutes, glass was allowed to return to room temperature, sandwiches were separated and placed in porcelain racks, and excess silane was removed by immersing in acetone. Coverslips were then dried quickly, avoiding condensation of water from the air, and then incubated with HO-PEG-NH2 by making “sandwiches” in petri dishes on a 75°C hot plate, in which the HO-PEG-NH2 powder melts. Air bubbles were pressed out carefully and PEG “sandwiches” were left overnight in a 75°C oven. Excess HO-PEG-NH2 was rinsed thoroughly using Milli-Q water and sonication in porcelain racks, and the passivated slides could then be dried and stored at 4°C between lens-cleaning tissue. Glass was generally used within 8 weeks of preparation.

#### Microscopy stock materials

Freshly prepared, ice-cold BRB80 buffer (80 mM K-PIPES, 1mM MgCl_2_, 1 mM EGTA, pH 6.8; prepared as a 5x stock and diluted to 1x before use) was used to resuspend β-casein (to 25 mg/mL), glucose oxidase (to 40 mg/mL), and catalase (to 20 mg/mL). Resuspended proteins were spun at 80,000 RPM in a TLA 100 rotor 4°C table-top ultra-centrifuge, aliquoted, snap-frozen in liquid nitrogen and stored at -80C. ATP and GTP were dissolved in MilliQ water to 100 mM each, adjusted to pH 7 using KOH, filtered with 0.22 μm pore size, and aliquots were stored at -80°C. Docetaxel was dissolved in DMSO to 1 mM, filtered with 0.22 μm pore size, and aliquots were stored at -80°C. Glucose was dissolved in MilliQ water to 1 M, filtered with 0.22 μm pore size, and stored at 4°C.

#### Flow chambers

On the day of experiments, flow cells were assembled from the PEG-passivated glass using 80 or 20 μm thick double-stick tape, generating a chamber about 5 mM wide and 7 mM long. With appropriately sized cover-glasses, multiple experiments could be performed in tandem. The assembled flow cells were kept for up to one day at room temperature and placed on a 33°C heat block just before preparing the assay.

#### Self-organization assay

On each day of experiments, BRB80 was freshly diluted from 5x stock (stored up to 2 weeks at 4°C) and kept on ice. HSET and KIF11 gel filtration buffer stocks without ATP or β-ME (stored at 4°C) were supplemented with ATP or β-ME and kept on ice (HSET gel filtration buffer: 50 mM Na-phosphate, 300 mM KCl, 1 mM MgCl_2_, 1 mM EGTA, 5 mM β-mercaptoethanol, 0.1 mM ATP, pH 7.5; KIF11 gel filtration buffer: 50 mM Na-phosphate, 300 mM KCl, 2 mM MgCl_2_, 10 mM β-mercaptoethanol, 0.1 mM ATP, pH 7.5). Aliquots of catalase and glucose oxidase were mixed and diluted with BRB80 to make the oxygen scavenger mix containing 18.1 mg/mL catalase and 11.8 mg/mL glucose oxidase. The oxygen scavengers were spun at 80,000 RPM in a TLA 100 rotor 4°C table-top ultracentrifuge, along with thawed β-casein to get rid of aggregates. The oxygen scavengers and β-casein were stored on ice.

Buffers and protein solutions were kept on ice, except for thawed docetaxel aliquots, which were kept at RT. Fresh KIF11 and HSET protein aliquots were thawed each experimental day, diluted appropriately in their respective gel filtration buffers. Labeled and unlabeled tubulin aliquots were thawed before each experiment, mixed for a final labelling ratio of 3.5% CF640R, and diluted appropriately in BRB80. The final experimental solution contained 19.2 μL tubulin mix, 5.76 μL KIF11 dilution or KIF11 gel filtration buffer, 5.76 μL HSET dilution or HSET gel filtration buffer, 2.12 μL β-casein, 2 μL oxygen scavenger mix, with 18.3 μL “docetaxel mix” to bring final buffer concentrations to 35 mM PIPES, 65 mM KCl, 1.68 mM MgCl_2_, 3.55 mM β-mercaptoethanol, 1 mM EGTA, 1 mM ATP, 0.64 mM GTP, 35 mM glucose, and 1 μM docetaxel (added freshly before each experiment from DMSO stock).

The sample was clarified by spinning at 13.3K RPM in a 4°C table-top centrifuge for 5 minutes. The supernatant was transferred to a new tube on ice. The prewarmed flow-cell was washed by flowing 50 μL of a “wash” solution with the same components as the final experiment solution, but KIF11, HSET, tubulin or oxygen scavengers were replaced by their respective storage buffers. The wash solution was replaced by flowing through 50 μL of the sample solution, using blotting paper to draw solution through the chamber. The chamber was sealed with silicone vacuum grease, transferred to an inverted spinning-disk confocal microscope with an incubator at 33°C. After locating the center of each flow cell (between the tape and glass boundaries), imaging was initiated between 3 and 4 minutes after initial temperature shift. Exposure times for each laser line were 200-300 ms, with images taken at intervals of one minute.

#### Polarity-marked microtubules

Stable polarity-marked microtubules, with a bright minus end and a dim plus end, were prepared using AlexaFluor568 tubulin. Polarity labelling was performed as previously described^43^. First, bright GMPCPP seeds were polymerized from 15 μM tubulin, with a total labelling ratio of 0.14, with 0.5 mM GMPCPP, diluted with BRB80 for 45 μL total volume, for 30 minutes at 37°C. The seeds were spun down and washed twice, and resuspended with 45 μL BRB80 at RT. 45 μL of polar extension mix containing 15 μM total of tubulin, of which 6 μM was NEM-tubulin, with AlexaFluor568 tubulin for a labelling ratio of 0.035, was mixed on ice with 1 mM GTP and 2 mM DTT. The mixture was transferred to 37°C for 1 min before adding 10% its volume of bright GMPCPP seeds and mixing gently, incubating for a further 20 minutes at 37°C. The polymerized extensions were subsequently stabilized by addition of docetaxel in two steps. After the 20 minute-polymerization, 5 μL BRB80 containing 2 mM DTT and 10 μM docetaxel was added. After 2 more minutes at 37°C, the seeds were diluted with 150 μL BRB80 with 2 mM DTT and 10 μM docetaxel. To remove the excess unpolymerized tubulin, seeds were spun and resuspended 3 times over a 0.22 μm spin filter, in BRB80 with 2 mM DTT and 10 μM docetaxel, for a final volume of 50 μL. Seeds could be stored at RT for several hours before significant end-to-end annealing. Seeds could be further diluted in BRB80 with 2 mM DTT and 10 μM docetaxel and sandwiched between untreated coverslips to assess efficiency of polarity marking. Of microtubules containing one bright segment, 63% had one dim segment (N = 747).

Microtubule gliding assays were performed as described previously^6^. Flow chambers were prepared using untreated glass coverslips and 70 μm double stick tape. The flow cells were washed with 50 μL BRB80, followed by 50 μL of the assay “wash mix”, and incubated 2 minutes for β-casein to assemble on the surface. 50 μL of 300 nM mGFP-KIF11 or avitag-mBFP-HSET in the self-organization assay buffer (without tubulin or oxygen scavengers) was added to the chamber and allowed to adhere to the surface for 5 minutes before washing out with two 50 μL washes of “wash mix”. Polarity-marked seeds were then diluted one in 200 in RT wash mix, and 50 μL of this polar seed sample was added to the chamber, which could then be sealed and examined on the confocal microscope. Motor attachment to the casein surface was confirmed by checking fluorescence of the appropriate motor fluorophore. Of mobile microtubules with one bright and one dim segment, 100% were moved in the bright (minus) direction in KIF11 gliding assays (N=35), and 97% in the dim (plus) direction in HSET gliding assays (N=36). Additionally, microtubules with a repeating bright-dim segment pattern were also moved towards the bright end by KIF11, indicating that annealed polarity-marked seeds retained directionality.

When polarity-marked seeds were to be added to active microtubule networks, they were first diluted 1 in 10 in BRB80 (to 1 μM docetaxel). The experimental solution for polarity-marked seed embedded networks contained 14.2 μL tubulin mix, 5,76 μL KIF11 dilution, 5.76 μL HSET dilution or HSET gel-filtration buffer, 2.12 μL β-casein, 2 μL oxygen scavengers, and 18.3 μL docetaxel mix on ice. The seeds were further diluted 1 in 3 in RT BRB80 directly prior to mixing the final sample to avoid addition of excess docetaxel while minimizing depolymerization. After spinning and before being added to the pre-warmed, washed flow cell, the solution was transferred into a RT tube, supplemented with 4.7 μL of the dilute polarity marked seeds, and mixed gently before flowing into the flow-cell and imaging as above.

### II. Simulations of active networks in Cytosim

Simulations were performed with Cytosim (https://gitlab.com/f-nedelec/cytosim), an Open Source project. Cytosim solves the Langevin equation of motion of bendable microtubules with crosslinkers in viscous medium^30^. Model parameters (see Table S1) were chosen near the experimentally determined values or, when experimental data were missing, chosen to best match the network behavior observed experimentally in this study. Simulation configuration files are included in Data S1, to allow reproducibility.

The system was simulated in 3D with X and Y dimensions of 20 µm and a thickness in Z of 0.2 µm unless otherwise stated. The thickness of the box was chosen to minimize computational cost while still allowing multiple filaments to cross each other when overlapped. The box is reflective in Z and periodic in X and Y. When microtubule densities are stated in the text, they are reported as the number of microtubules in the quasi-2-dimensional box per X-Y area. Simulations were run with a time step of 0.005 s for a duration of 8000 s (∼2h), corresponding to experimental time scales.

#### Microtubule model

Microtubules interact with each other via Hookean soft-core repulsive forces directed perpendicular to the filament axis so as to minimize sliding force parallel to the microtubule axis. Microtubules have discrete lattice binding sites covering 8 nm to which the crosslinker units can bind. Microtubules were nucleated from randomly distributed “seeds” at a rate such that most microtubules were nucleated at the very beginning of the simulation, similar to the experiment. Once nucleated, the plus end of the microtubule grows at a gradually decreasing speed, to mimic the limited availability of soluble tubulin pool as in the experiments (Suppl. Fig. 1). The time-dependent growth rate is expressed as *V*_*g*_ (*t*) = *α*[1 − {∑ *L*_*i*_ (*t*)}/Ω] where *α* is the maximum growth speed, ∑ *L*_*i*_(*t*) is the total length of all microtubules at time t, and Ω is the available amount of tubulin subunits in the system, a parameter expressed in µm.

To mimic the broadly distributed microtubule lengths in the experiment, we introduced variability in growth speed for each microtubule. We assigned a random speed *α* drawn from a normal distribution *N(0,1)* as *α* = *α*_0_ |*1+0*.*25 N(0,1)*|. The resulting speed distribution is centered near *α*_0_ and restricted to 0—2 µm/s. Fully grown microtubules did not undergo catastrophe, mimicking the docetaxol-stabilized microtubules in the experiment. At steady state, the average microtubule length is 2.5 µm, and thus shorter compared to the experiment. This was a decision made necessary to keep the simulation running time manageable on the computing cluster, in particular for large systems of which their dimensions exceed the individual microtubule lengths many times.

#### KIF11 model

KIF11 was modelled as a symmetrical crosslinker with a pair of motor units connected by a Hookean spring with non-zero rest length. A freely diffusive motor in the medium can bind to a microtubule at a constant rate if it is within a specified binding distance. A bound motor can unbind with a force-dependent rate. Once bound, the motor walks stochastically on the 1D lattice toward the plus end in a processive manner at a speed linearly dependent on the load. The velocity of a motor varies with force 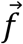, as 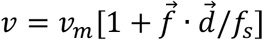, where 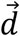 is the direction in which the motor would move along the filament if it was unloaded, *f*_*s*_ is the stall force and *v*_*m*_ is the unloaded motor speed. Speed is thus decreased by antagonistic force as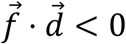. KIF11 can crosslink a pair of microtubules with both units but cannot bind with both units to the same microtubule. The force is Hookean with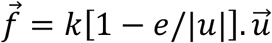, for a spring of rest length *e*, stiffness *k* and extension 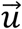. The joints formed by the link rotate freely such that the angle between the crosslinked microtubules is not constrained.

As shown in previous simulations^6^, microtubules organize into a nematic network when the steady-state motor profile along the microtubule is uniform. However, with short microtubules that stop growing and do not undergo catastrophe, KIF11 motors are able to reach and accumulate at the microtubule plus end, obstructing nematic network formation. This can be resolved by having longer microtubules in a larger simulation box, but this would incur a prohibitive computational cost. Instead, we set the motors to unbind immediately at the end of the microtubule, which allows nematic network organization with short microtubules and thus at reasonable computing time.

#### HSET model

HSET was modeled as an asymmetric crosslinker with one motor unit and one diffusive unit. The motor is minus-end directed and non-processive. When bound, the diffusive subunit steps stochastically to neighboring lattice sites on the microtubule. When the motor is not bound, the force is null and the diffusible subunit undergoes unbiased diffusion. However, when the motor is bound, the force would bias the plus- and minus-end directed hopping rates in a thermodynamically consistent manner, as described previously^44, 45^. Unlike KIF11 motor units, the HSET motor unit only binds transiently to a microtubule, taking on average 2 steps before unbinding. In comparison, the diffusive unit dwells for a relatively longer time on the microtubule. When reaching the end of a microtubule, both units can stay bound, allowing clustering of microtubule ends.

#### Quantification of microtubule motion

To calculate the overall speed of microtubule motion in Fig. 5 and Fig. 6, we extracted the positions of microtubule minus ends at a time interval Δt. We then calculated the displacement component parallel to the microtubule axis. Averaging the displacement for all microtubules and dividing by Δt gave the overall averaged microtubule speed. A large Δt (640 or 1280 s) was chosen such that the displacement due to diffusion is negligible relative to directed motion. The overall speed approaches the motor’s speed when all microtubules are crosslinked and slid continuously. In the absence of motor, the overall averaged speed as extracted here has a small, non-zero value due to the rotational diffusion of the microtubules. As a matter of convention, a positive speed indicates minus-end leading unidirectional sliding (plus-motor driven). Likewise, a negative sign indicates sliding with the plus-end leading.

#### Bundle and aster size calculation

To calculate the size of bundles and asters as shown in Fig. 5c, we first identify clusters of microtubules crosslinked by the same type of crosslinker. The number of microtubules in the largest cluster are then reported as the bundle size and the aster size respectively for KIF11 and HSET. The size was normalized by the number of microtubules for convenience.

## ACKNOWLEDGEMENTS

We thank Johanna Roostalu for designing and cloning the biotinylated HSET construct. We thank Claire Thomas and Raquel Garcia-Castellanos for biochemistry support and Davide Normanno for microscopy support. Simulation were performed on the high-performance computing cluster at the Centre for Genomic Regulation (CRG) in Barcelona, Spain. This work was supported by the Spanish Ministry of Economy, Industry and Competitiveness to the CRG-EMBL partnership, the Centro de Excelencia Severo Ochoa and the CERCA Programme of the Generalitat de Catalunya, and the Francis Crick Institute, which receives its core funding from Cancer Research UK (FC001163), the UK Medical Research Council (FC001163), and the Wellcome Trust (FC001163). W.-X. C. is supported by an HFSP fellowship (LT000682/2020-C). F.N. is supported by the Gatsby Charitable Foundation (Grant PTAG-024) and the European Research Council (ERC Synergy Grant, project 951430). T.S. acknowledges support from the European Research Council (Advanced Grant, project 323042, ERC Synergy Grant, project 951430).

---

**Table S1.**

| Parameter | Value | Note |
| --- | --- | --- |
| **Simulation** |  |  |
| Time step | 5 ms | Small enough for convergence |
| Viscosity | 0.02 pN s/$\mu\text{m}^2$ | Close to water |
| Box thickness | 0.2 $\mu\text{m}$ | Thick enough for 3-4 filaments to cross |
| **Microtubule** |  |  |
| Rigidity | 30 pN $\mu\text{m}^2$ | Ref. 46 |
| Steric radius | 0.05 $\mu\text{m}$ | Ref. 47 |
| Steric force constant | 50 pN /$\mu\text{m}$ | Constrained by thermal fluctuations and force of motors |
| Average length | 2.5 $\mu\text{m}$ | Optimum for computation |
| Growth rate | 0.03 $\mu\text{m/s}$ | Ref. 6. |
| Sensitivity of growth to force | 1.67 pN | Ref. 46 |
| Nucleation rate | 0.1 /s | Time for all microtubules to nucleate is short relative to total simulation time |
| **KIF11 motor** |  |  |
| Stall force | 5 pN | Ref. 48 |
| Unbinding force | 5 pN | Refs. 49, 50 |
| Maximum motor speed | 0.03 $\mu\text{m/s}$ | Ref. 6 |
| Binding rate | 0.5 /s | Estimated to match experiment |
| Unbinding rate | 0.1 /s | From run length =0.3 $\mu\text{m}$ as measured in Ref. 20 |
| Binding radius | 0.16 $\mu\text{m}$ | 1.5x crosslinker rest length |
| **HSET motor** |  |  |
| Stall force | 5 pN | Same as KIF11 |
| Unbinding force | 5 pN | Same as KIF11 |
| Maximum motor speed | - 0.08 $\mu\text{m/s}$ | Within measured range in Refs. 6, 10, 32 |
| Binding rate | 5 /s | Estimated to match experiment |
| Unbinding rate | 5 /s | Same order of magnitude as in Ref. 24 |
| Binding radius | 0.16 $\mu\text{m}$ | 1.5x crosslinker rest length |
| **HSET Diffusive tail** |  |  |
| Diffusion constant | 0.1 $\mu\text{m}^2/\text{s}$ | Refs. 10, 24 |
| Binding rate | 0.5 /s | Same as KIF11 motor |
| Unbinding rate | 0.01 /s | Ref. 24 |
| Unbinding force | 5 pN | Same as motor domain |
| Binding radius | 0.16 $\mu\text{m}$ | 1.5x crosslinker rest length |
| **Motor complex** |  |  |
| Rest length | 0.105 $\mu\text{m}$ | 2x microtubule radius (0.025 $\mu\text{m}$) + actual crosslinker rest length (0.08 $\mu\text{m}$) |
| Link stiffness | 100 pN/$\mu\text{m}$ | Ref. 51 |
| **Discretization parameter** |  |  |
| Filament segment length | 0.5 $\mu\text{m}$ | Optimum for computation |
| Filament lattice size | 8 nm | Size of a tubulin dimer |
| Motor stepping size | 8 nm | Same as lattice size |

## SUPPLEMENTARY FIGURE LEGENDS

**Suppl. Fig. 1.**
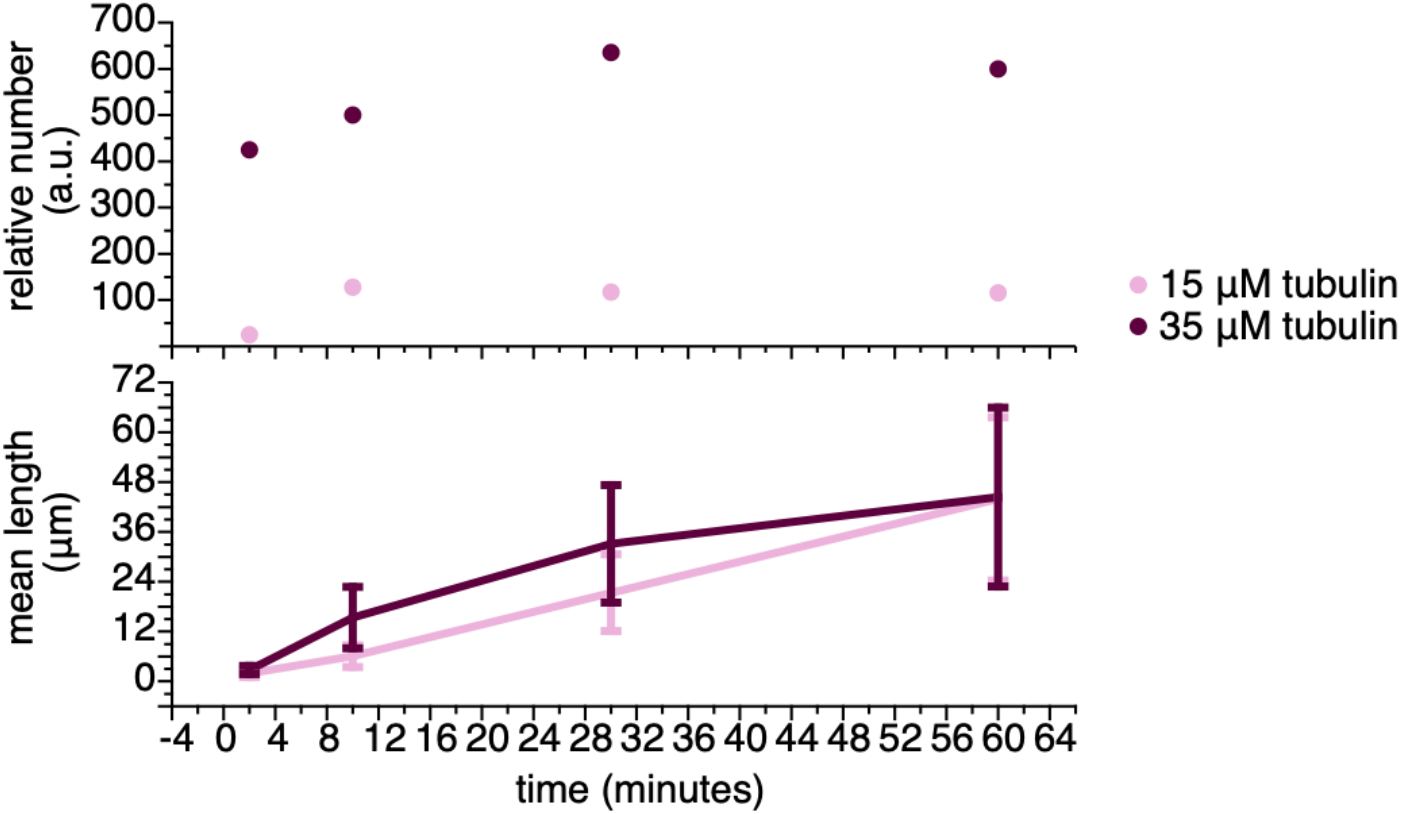
Plots of relative number (above) and mean length (below) at different time points, taken from microscopy images of diluted samples immobilized on a rigor-mutant kinesin surface (surface prepared as published ^41^. The samples for 35 μM tubulin were diluted five times more, so that the relative number reports five times the counted number in the field of view. Error bars on the mean length are standard deviations. For time points at 2, 10, 30, and 60 minutes the number of microtubules used to calculate the mean lengths were 25, 128, 117, and 116 (15 μM tubulin); 85, 100, 127, and 120 (35 μM tubulin).

**Suppl. Fig. 2.**
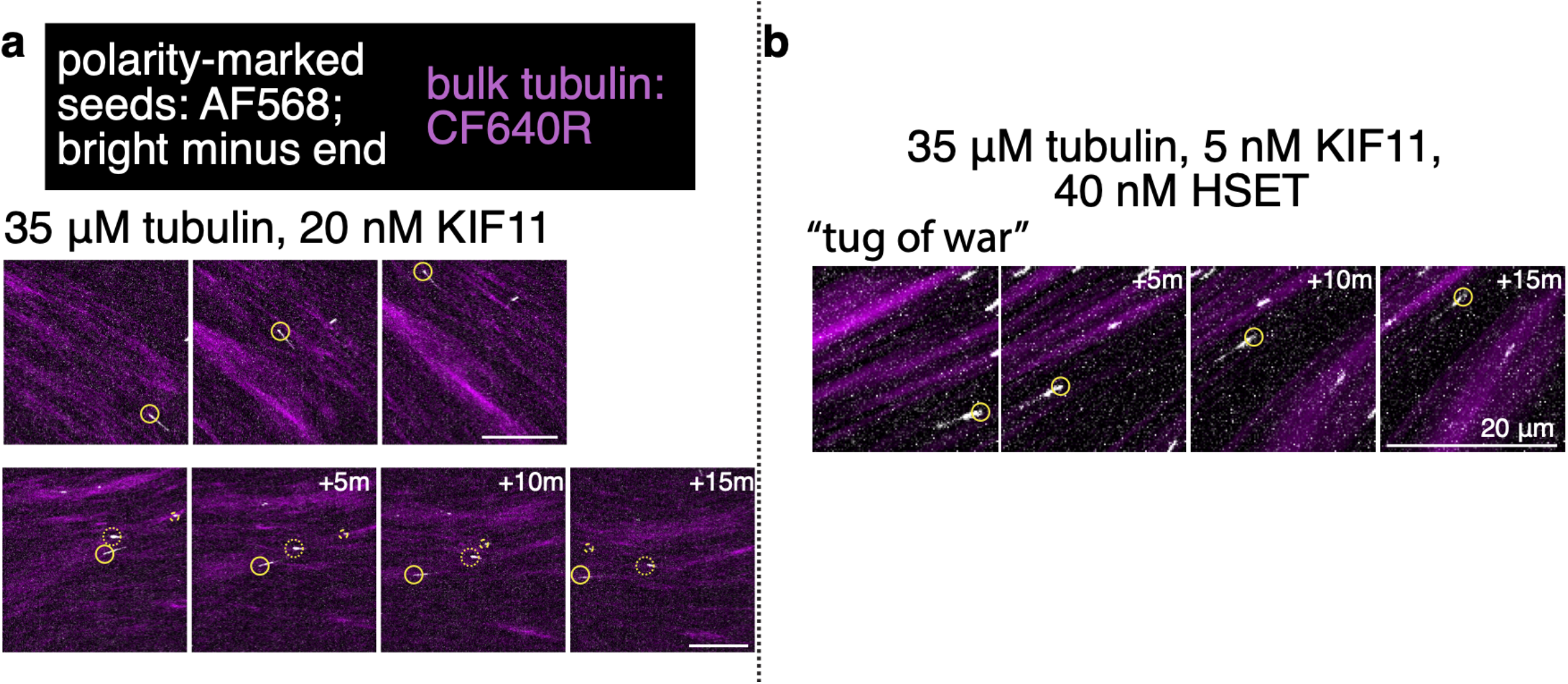
Motility of polarity-marked microtubules with bright minus ends (white; AlexaFluor567) in high density microtubule networks (bulk tubulin, purple; CF640R). Yellow circles indicate microtubule minus end. a 35 μM tubulin, 20 nM KIF11-mGFP: microtubules moving towards their minus ends. b 35 μM tubulin, 40 nM mBFP-HSET, 5 nM KIF11-mGFP: microtubule undergoing saltatory, “tug of war” behavior.

**Suppl. Fig. 3.**
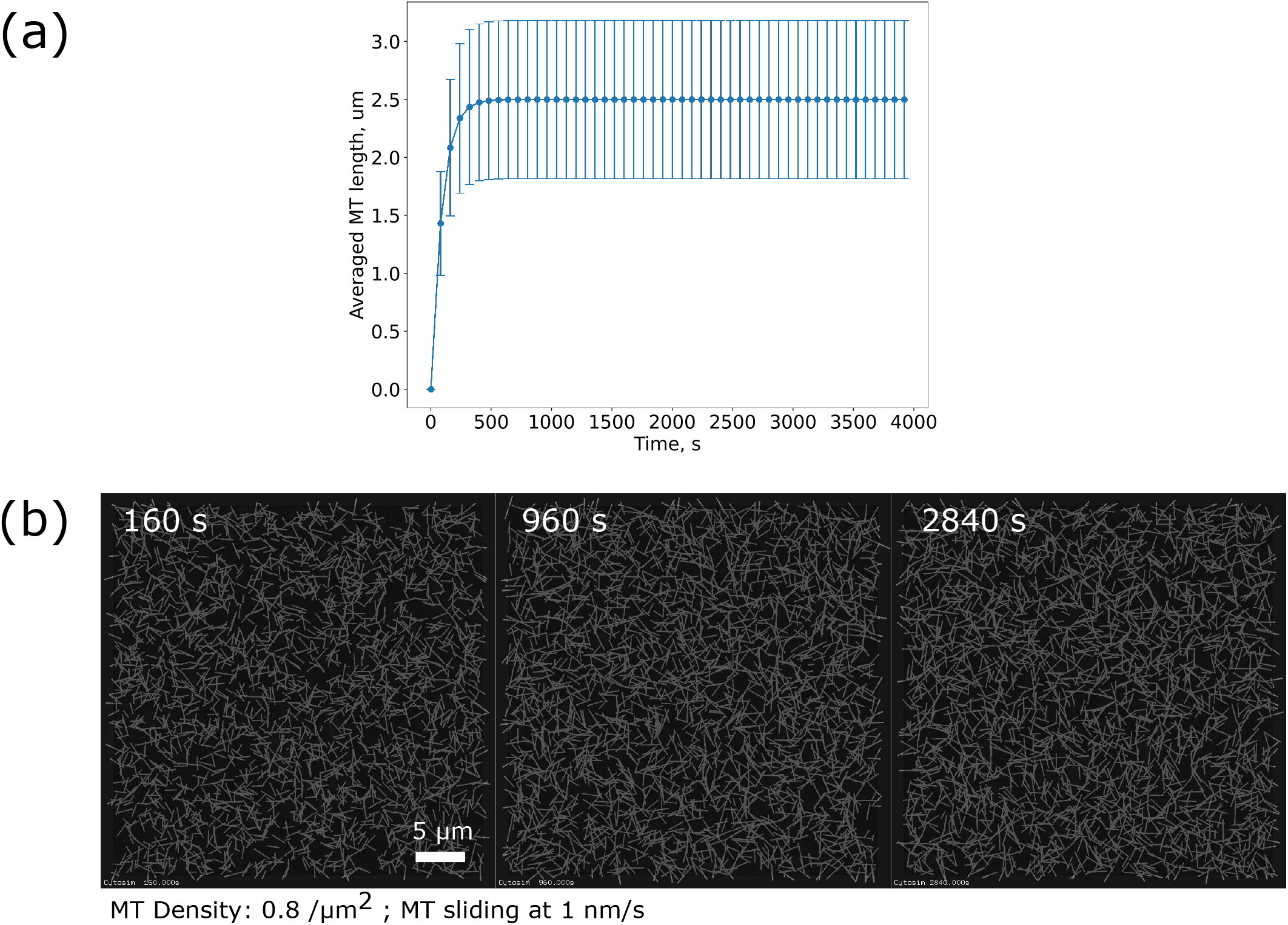
Simulated microtubule nucleation and growth. **a**, Time series of the average length of 2880 microtubules up to 4000 s, from a simulation covering 60×60×0.2 µm. **b**, Corresponding time evolution of random microtubule organization in the absence of crosslinkers.

**Suppl. Fig. 4.**
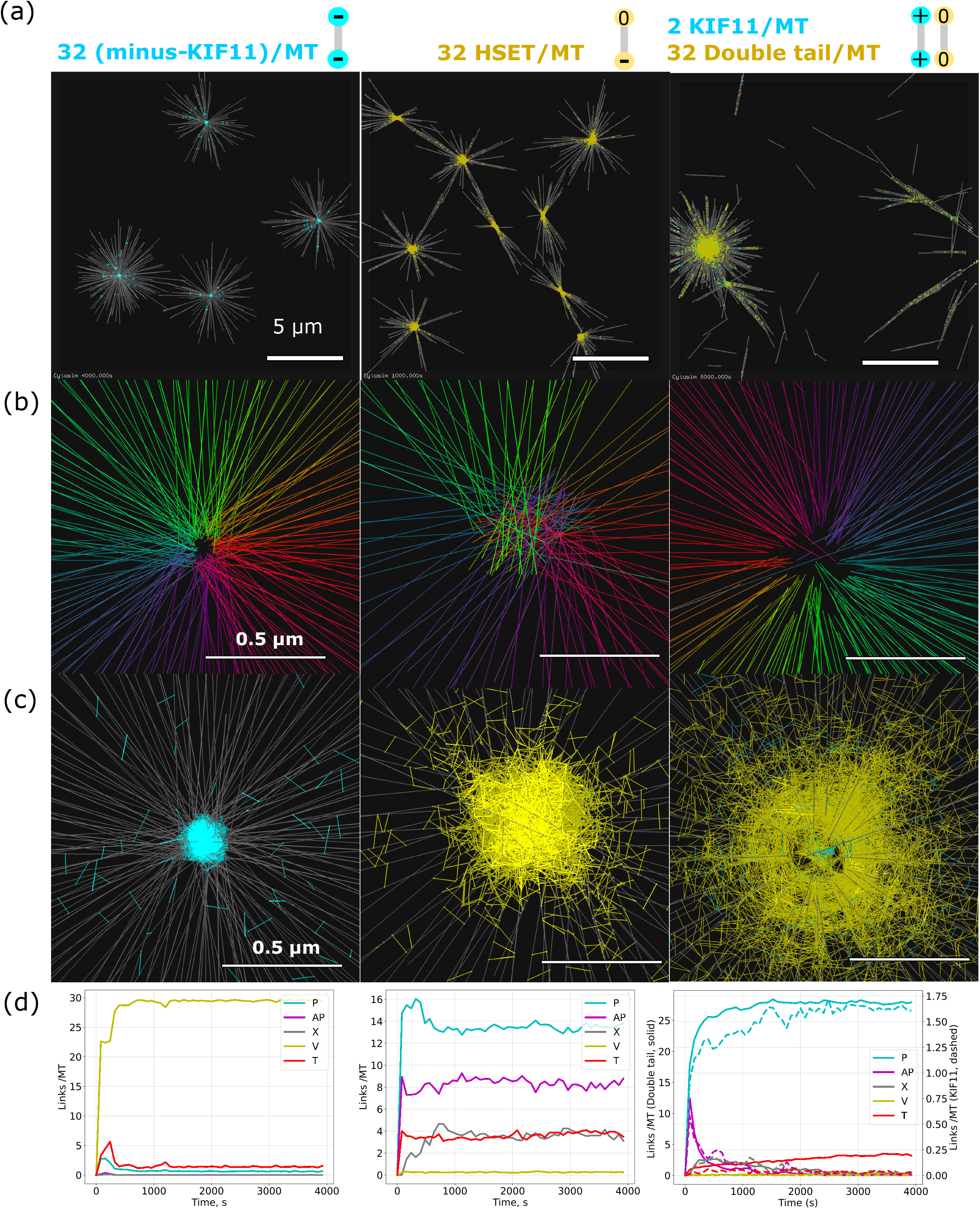
Aster formed by different crosslinker types. **a**, Asters formed by KIF11 with hypothetical minus-end directed motors (left), by HSET (middle) and by plus-directed KIF11 mixed with double tail HSET variant (right). **b**, Close-up view of aster’s center with microtubules colored according to its orientation. Microtubules’ ends are focused (left), overlapped (middle) and focused (right) **c**, Localization of crosslinkers at the aster’s center: minus KIF11 are narrowly localized (left), HSET are broadly localized (middle); double tails are localized with a radial gradient and KIF11 are narrowly localized at the center (right). **d**, Time evolution of the connection types. Links made by crosslinkers were classified into 5 categories (V, T, P, AP, X) according to position of the crosslinker’s unit with respect to the microtubules’ ends, and the angle between the microtubules ^6^. V-links connect microtubules’ ends; T-links connect microtubules’ end to the side of another microtubule; P-links connects parallel microtubules where the internal angle is smaller than 60 degrees; AP-link connect antiparallel microtubules with an angle between 120 and 180 degrees; X-links connect microtubule’ sides when these microtubules form an angle from 60 to 120 degrees. The count of links was normalized by the number of microtubules. Left: a high number of V-links indicate that the ends of microtubules are connected into focused asters; Middle: a prominence of T-links over V-links indicate that the ends of microtubules are not sharply focused. Instead, a lot of P, AP and X-links suggest that microtubules overlap and are bundled; Right: many P-links and few V-links by KIF11 (dashed line) indicate KIF11 bind mostly on parallel microtubules and fall off when reaching the plus ends. Both P and T links by double tail (solid lines) are high, indicating that the double tail is helping the aster to stay focused.

**Suppl. Fig. 5.**
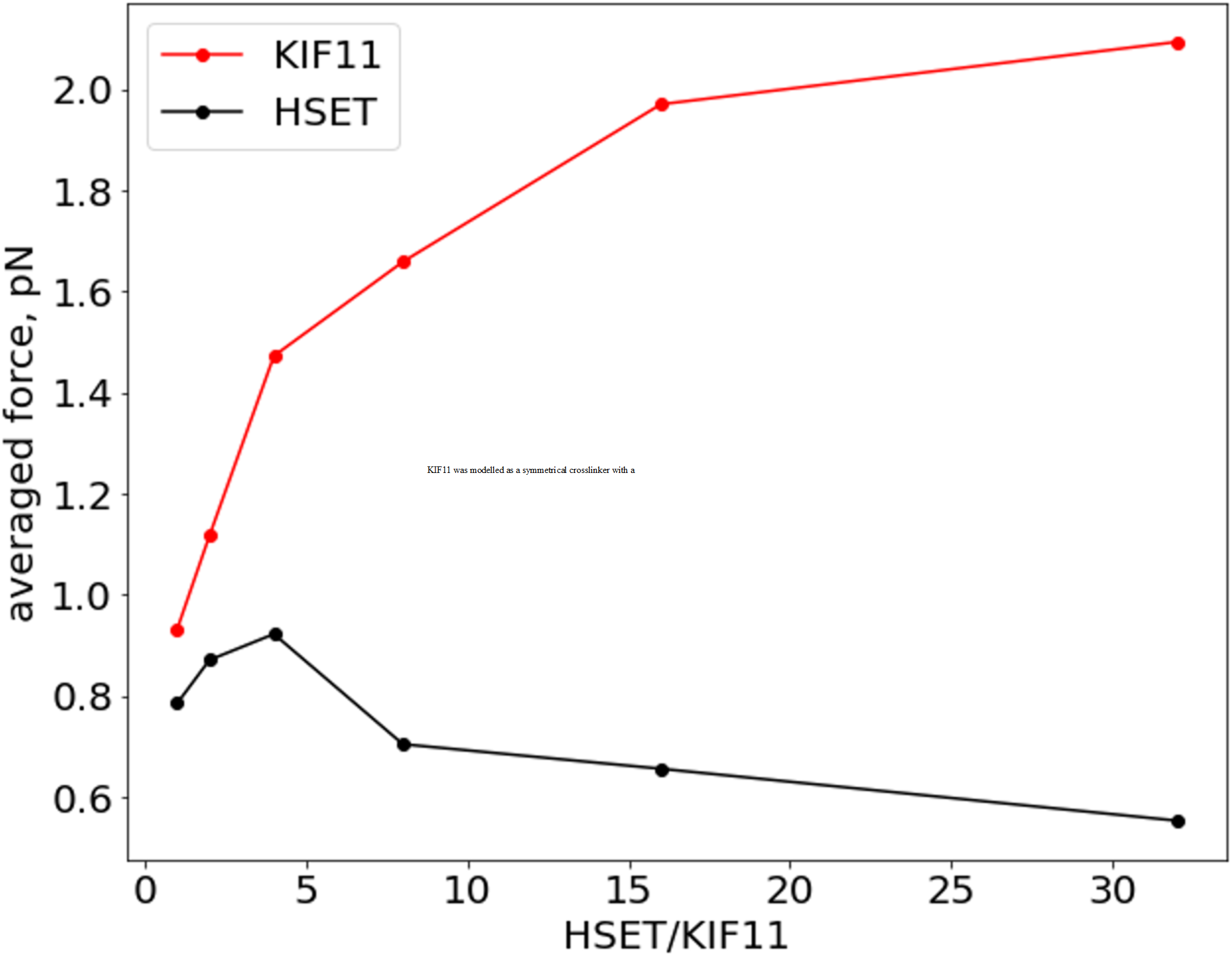
Tension of crosslinked KIF11 and HSET. Averaged force exerted by KIF11 (red) and HSET (black) when sliding antiparallel microtubules. The force is plotted against the ratio of the number of HSET and KIF11. Overall, the tension produced are lower with HSET compared to KIF11 due to its slippery unit. Tension of KIF11 increases when more HSET are introduced. Simulation contains 20 microtubules initialized in an antiparallel arrangement.

**Suppl. Fig. 6.**
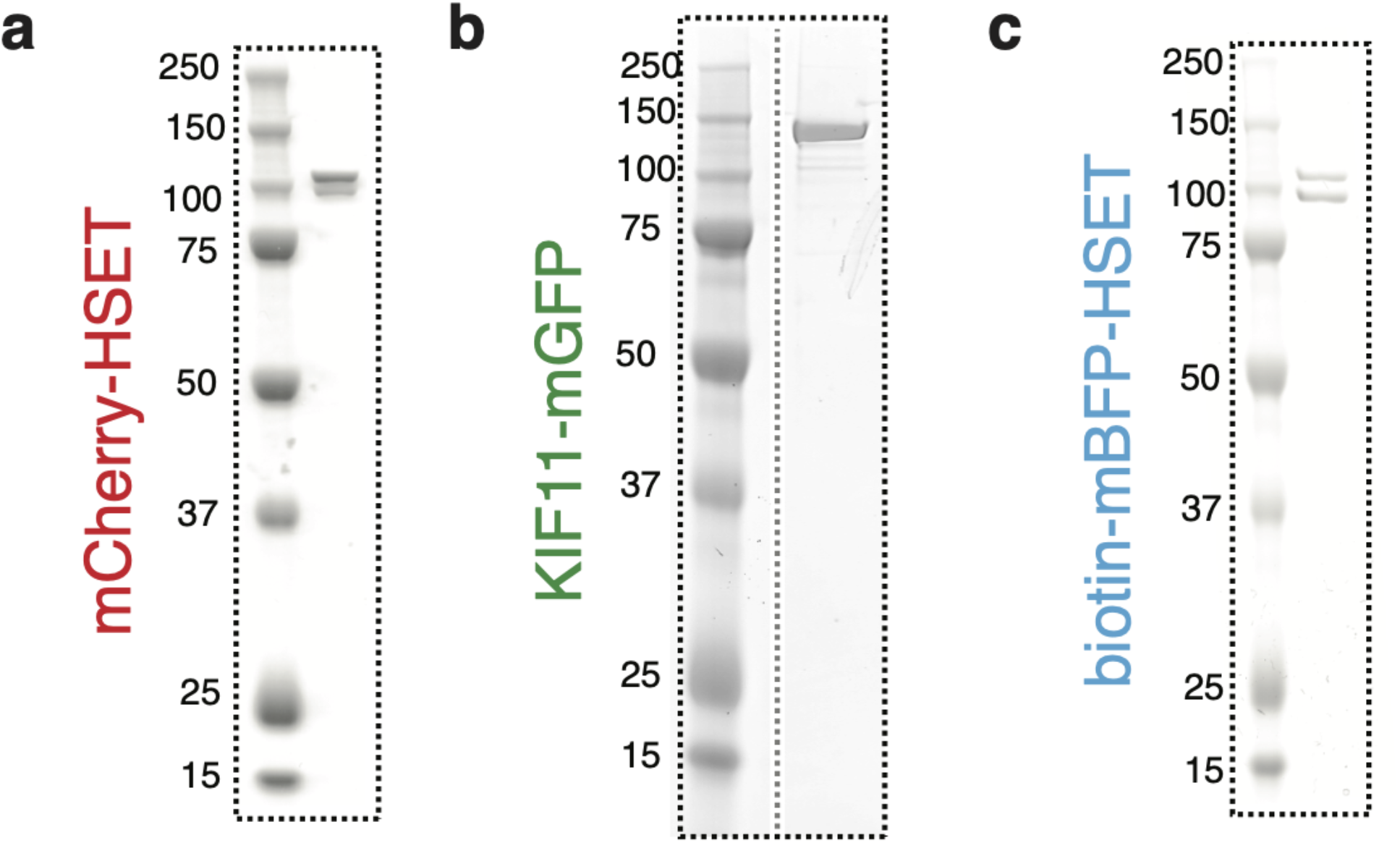
Coomassie-stained SDS-PAGE gels of the three recombinant motor proteins. **a** mCherry-HSET, **b** KIF11-mGFP, **c** biotin-mBFP-HSET. Gel images are cropped and rotated to align to dashed boxes. KIF11-mGFP panel stitches protein marker and purified protein lanes from the same gel, translated on the x axis.

## MOVIE LEGENDS

**Movie 1** | 35 µM tubulin networks driven by **a**, 10 nM HSET and **b**, 60 nM KIF11. Related to Fig. 1.

**Movie 2** | 10 µM tubulin networks driven by 20 nM HSET with **a**, 5 nM KIF11 and **b**, 20 nM KIF11. Related to Fig. 2.

**Movie 3** | 35 µM tubulin networks driven by 5 nM KIF11 and 0 nM HSET, 5 nM HSET, 40 nM HSET and 100 nM HSET. Related to Fig. 4a.

**Movie 4** | 15 µM tubulin networks driven by 5 nM KIF11 and 0 nM HSET, 5 nM HSET, 10 nM HSET and 40 nM HSET. Related to Fig. 4b.

**Movie 5** | Polarity-marked seeds with bright minus ends translocating in a mixed-motor nematic network formed from 35 µM tubulin, 40 nM HSET, and 5 nM KIF11. Minus ends manually tracked and labeled with circles. **a, b**, Seeds sliding towards their minus ends in bundles. **c**, Seed exhibiting saltatory behavior. Related to Fig. 4d.

**Movie 6** | **Left**, Movie of simulated KIF11-driven extensile nematic bundle in a thin 3D box (dimension: 60×60×0.2 µm). The 2880 microtubules are colored in gray, and the 46080 KIF11 in cyan. **Right**, Movie of simulated HSET-driven aster formation. Simulation space is 40×40×0.2 µm with 1280 microtubules and 40960 HSET. Related to Fig. 5.

**Movie 7** | **Left**, Random microtubule organization in the presence of low amount of KIF11 (cyan); **Middle**, Extensile bundle formed when HSET (yellow) was added in the presence of KIF11 at 2:1 ratio. **Right**, Minus aster formation when HSET and KIF11 were mixed at 16:1 ratio. Simulation space is 60×60×0.2 µm with 2280 Microtubules and 5750 KIF11. Related to Fig. 6a.

**Movie 8** | A portion of an extensile bundle formed in the presence of HSET and KIF11 at 2:1 ratio. **Left**, Microtubules (gray) with HSET (yellow) and KIF11 (cyan). **Right**, A few microtubules in the bundle are drawn as arrows (plus end = head), and colored randomly for this visualization. They move with the minus ends leading. Related to Fig. 6b.

